## Supplementary Figures for "Y chromosome sequence and epigenomic reconstruction across human populations"

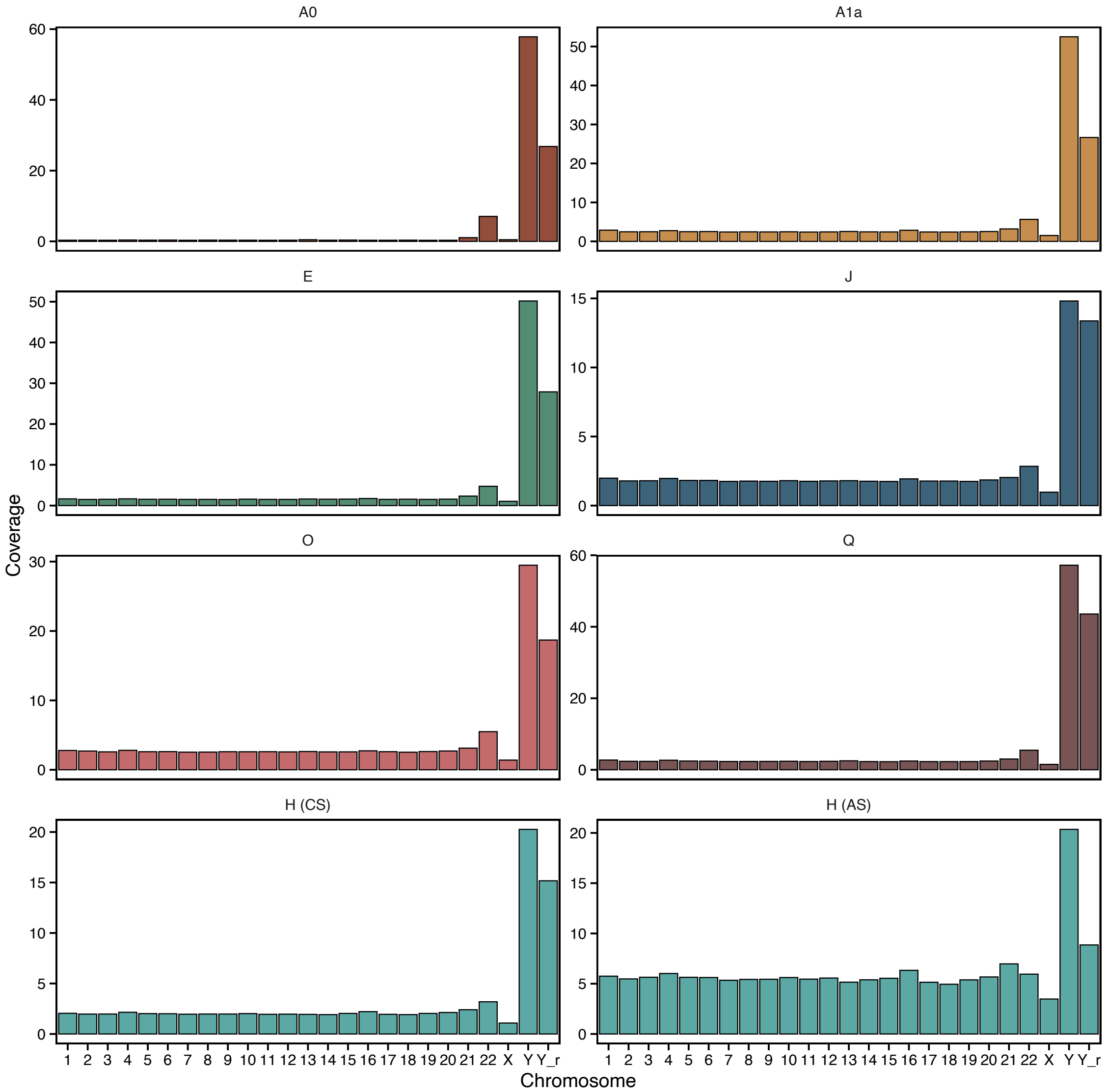


**Supplementary Figure 1.** Coverage of the 22 autosomal chromosomes, X chromosome, Y chromosome, and the random contig of the Y chromosome (Y_r) by haplogroup. H haplogroup separated by methodology: CS refers to chromosome sorting and AS refers to adaptive sampling.

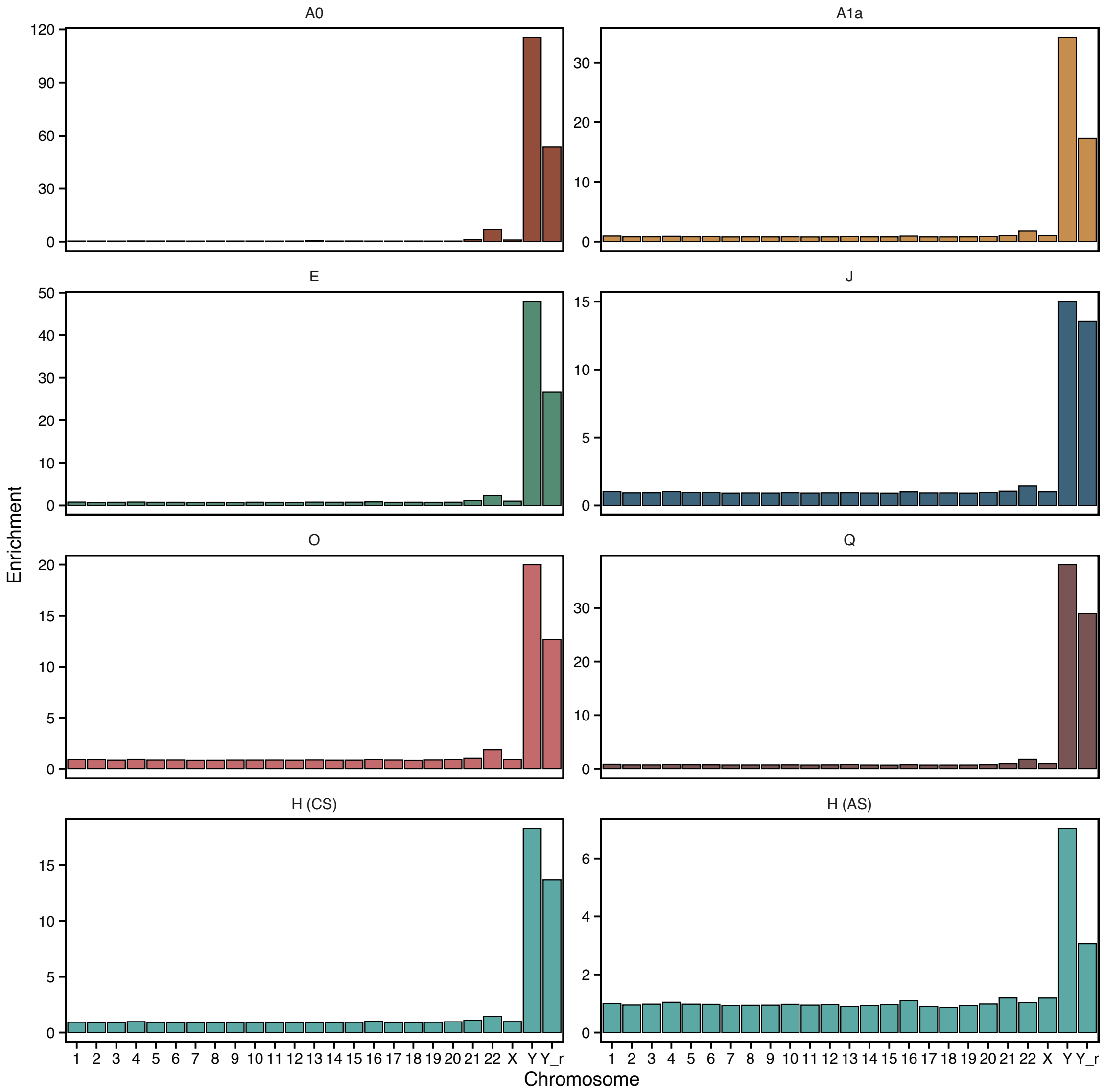


**Supplementary Figure 2.** Enrichment specificity of the sequencing data generated for the 22 autosomal chromosomes, X chromosome, Y chromosome and the random contig of the Y chromosome (Y_r) by haplogroup. H haplogroup separated by methodology: CS refers to chromosome sorting and AS refers to adaptive sampling.

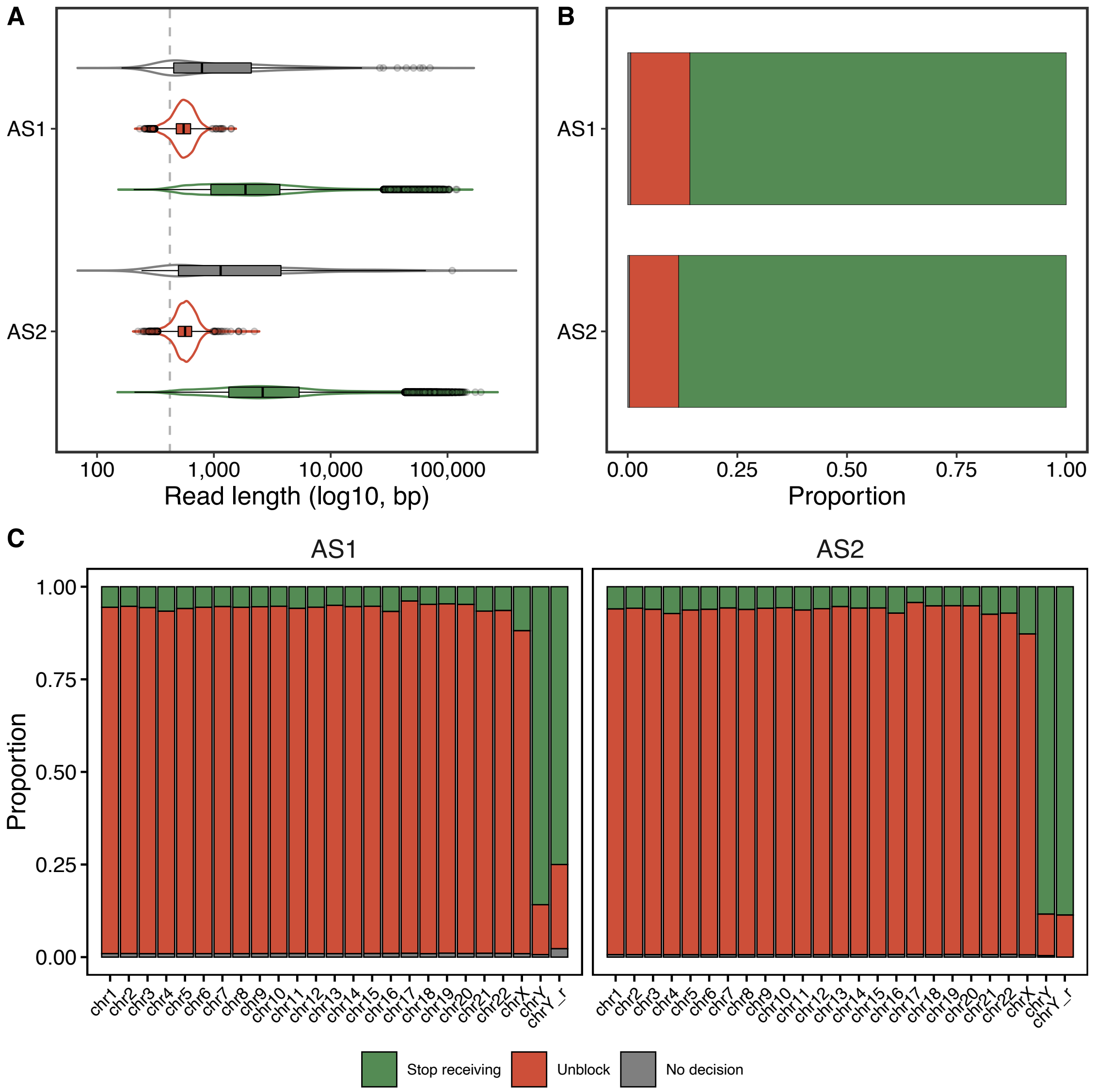


**Supplementary Figure 3.** Adaptive sampling (AS) metrics in the two runs generated for the H haplogroup sample (AS1 and AS2). (A) Read length and (B) Proportion of reads mapping to the GRCh38 chromosome Y (chrY) or the random chromosome Y contig (chrY_r) grouped by AS read decision in each AS run. (C) Chromosome-wise proportion of mapping reads in each AS run grouped by read decision type. The dashed grey line in (A) indicates 420 bases, the theoretical length each read is sequenced before the sequencer makes a decision. AS decision types: *unblock* reads are rejected by the pore, *no decision* reads are inconclusive, and *stop receiving* reads are further sequenced.

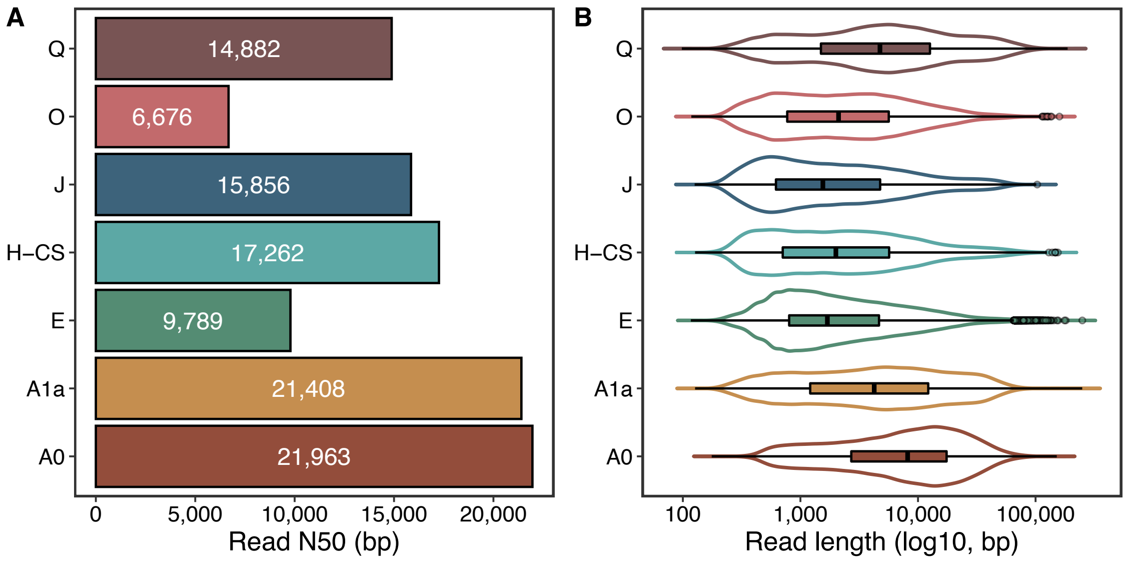


**Supplementary Figure 4.** (A) Read length N50 values. (B) Read length distributions of reads mapping to the GRCh38 chromosome Y or the random chromosome Y contig for each haplogroup. In (A) read lengths are shown in log10 scale. Only the chromosome sorting data for the H haplogroup is shown (H-CS).

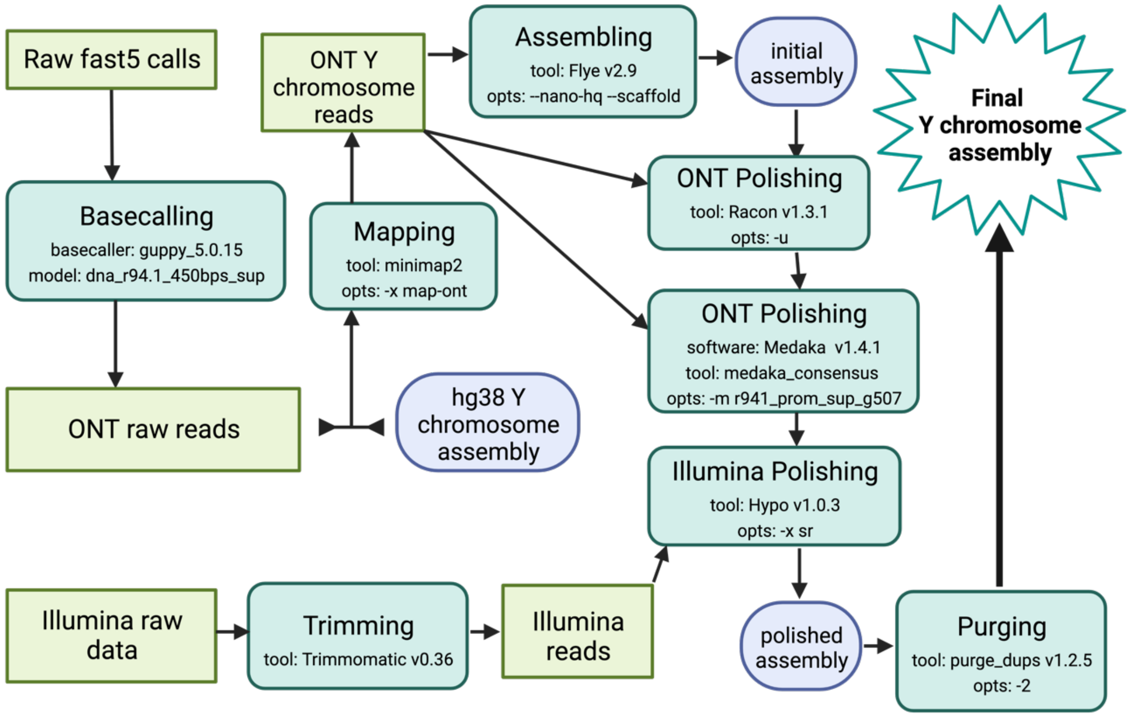


**Supplementary Figure 5.** Reference-guided Y chromosome assembly pipeline using chromosome sorting data. We consider both chrY and random chrY contig as the hg38 Y chromosome assembly. opts = running options; ONT = Oxford Nanopore Technologies. Created with BioRender.com.

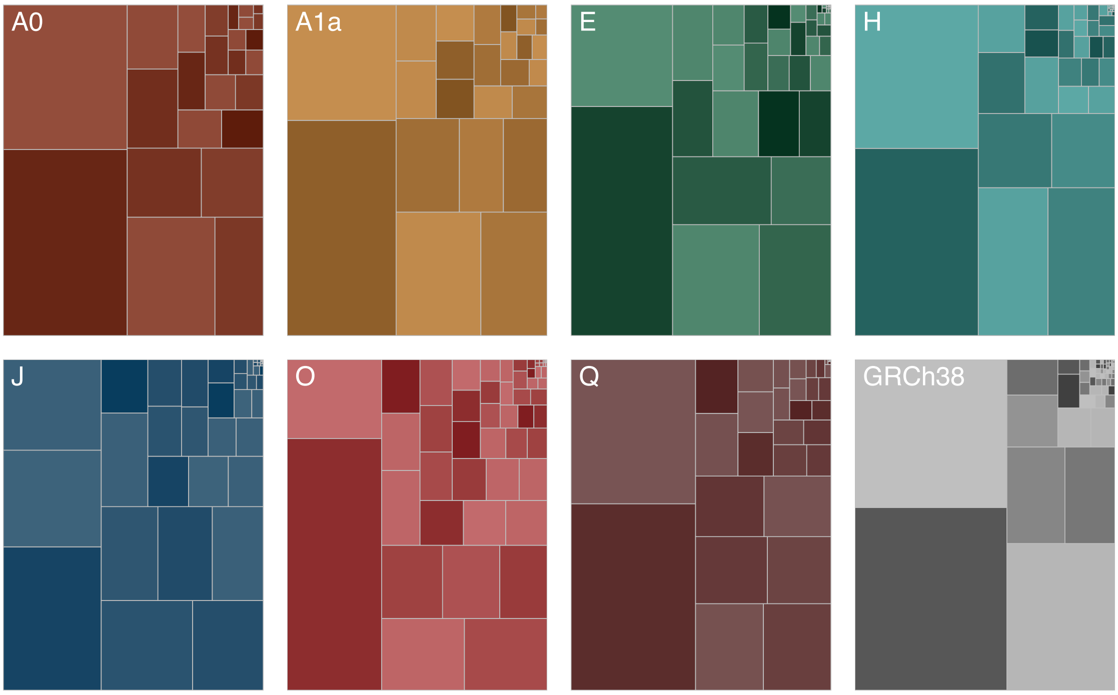


**Supplementary Figure 6.** Treemaps comparing the contiguity of the assemblies obtained of the different chrY haplogroups and GRCh38 chrY. The size of each rectangle corresponds to the size of a contig within each of the assemblies.

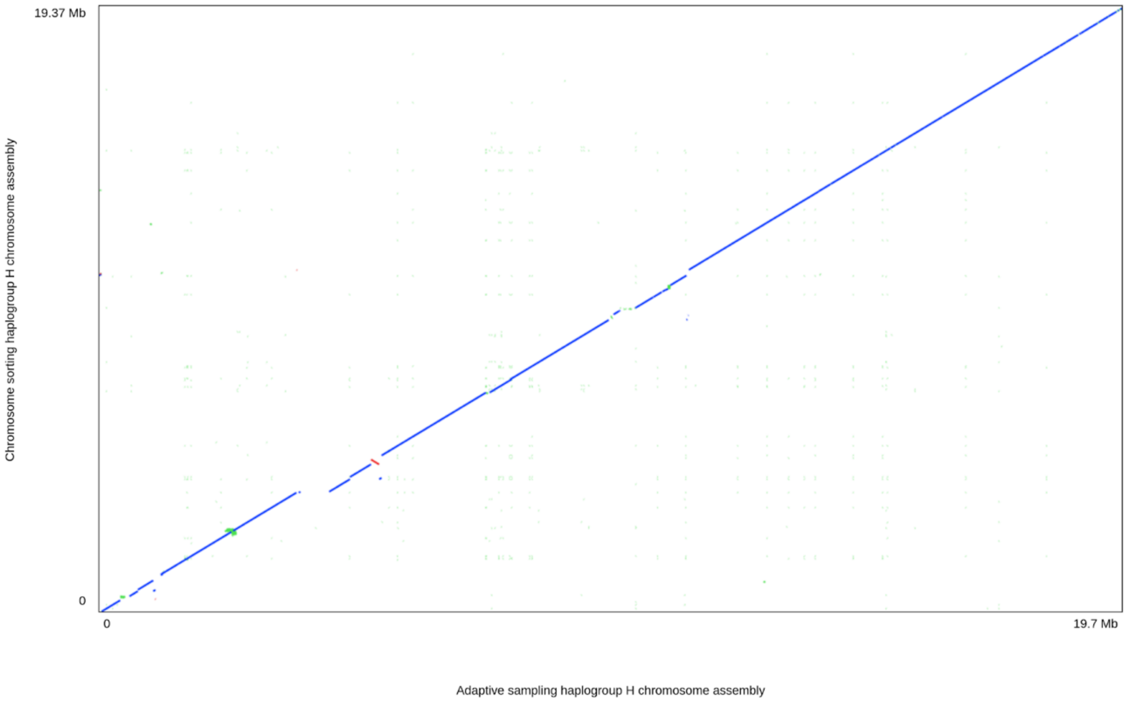


**Supplementary Figure 7.** Dot-plot comparing haplogroup H (cell line GM21113) assemblies, manually scaffolded based on alignments to GRCh38, obtained using same amount of data of two different selective sequencing methods. The assembly generated from adaptive sampling is shown in the X axis and the assembly generated from chromosome sorting data is shown in the Y axis.

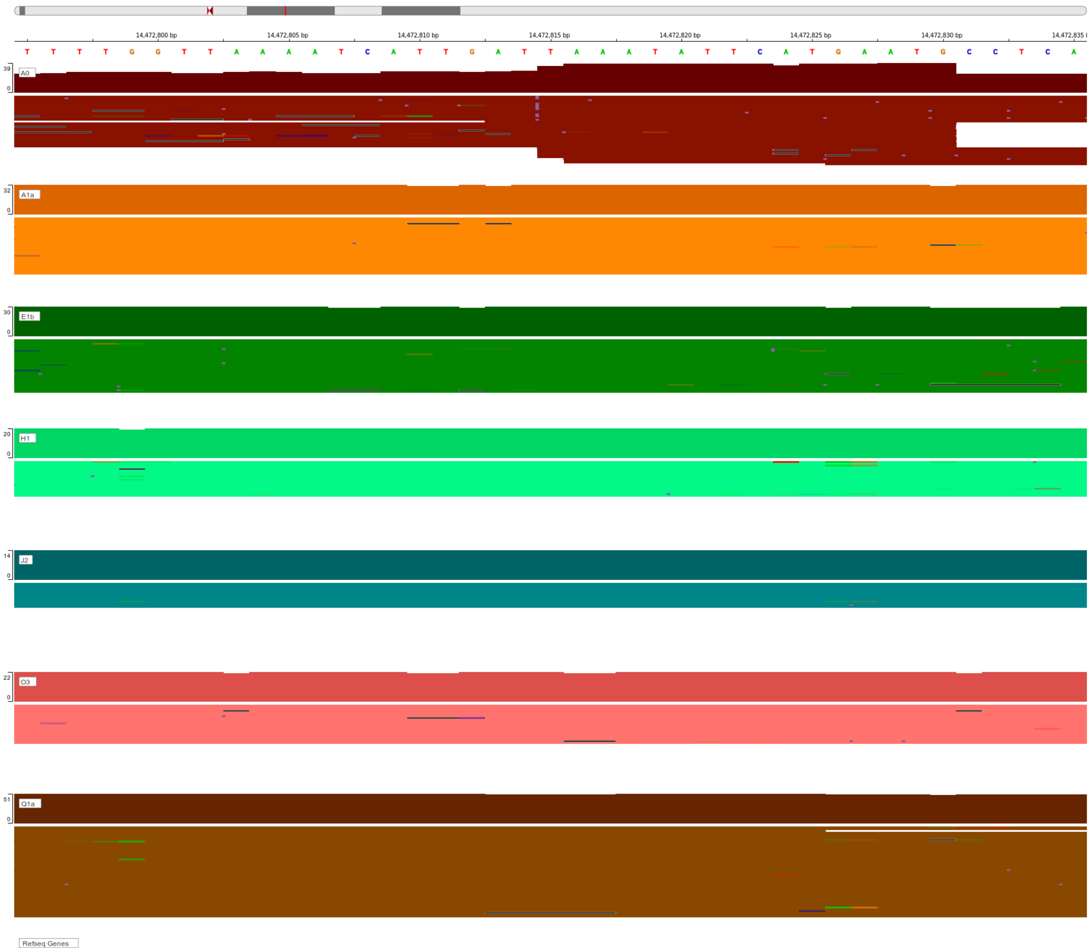


**Supplementary Figure 8.** IGV-screenshot showing ONT reads of 7 different Y haplogroups mapping to GRCh38 region with the longest insertion (shown in violet) found using *Sniffles.* It is only detected in the A0 haplogroup reads and not in the rest.

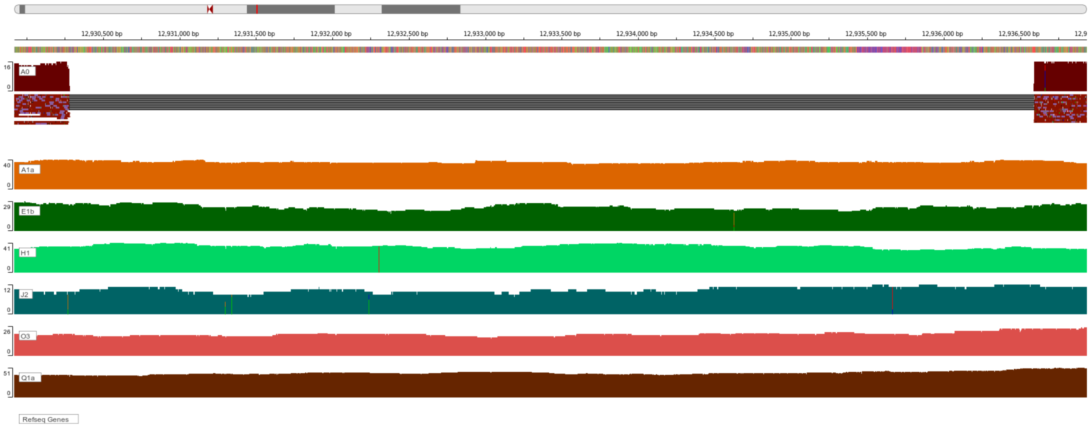


**Supplementary Figure 9.** IGV-screenshot showing ONT reads of 7 different Y haplogroups mapping to GRCh38 region with the longest deletion found using *Sniffles.* It is observed in the A0 haplogroup.

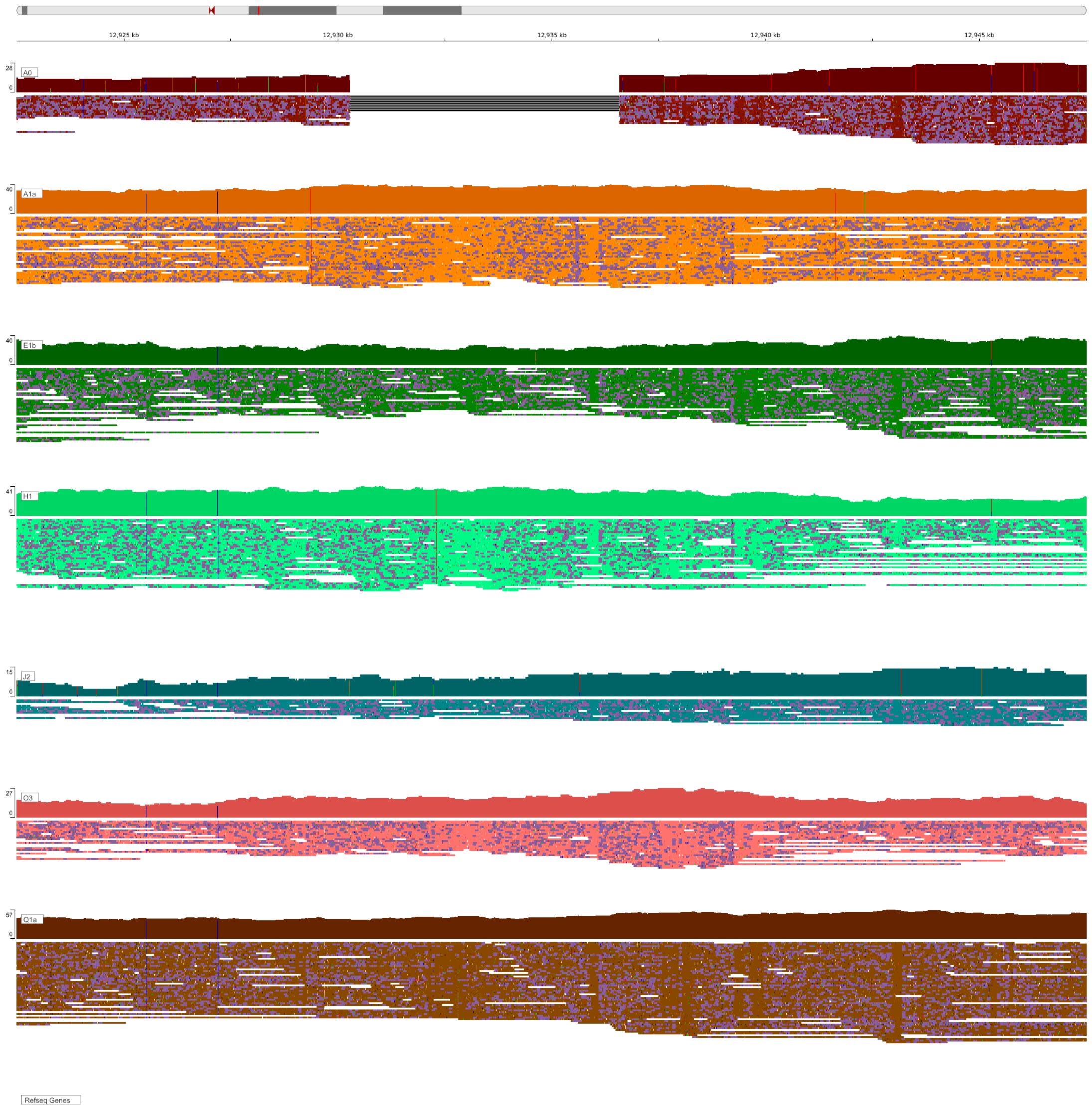


**Supplementary Figure 10.** IGV-screenshot showing ONT reads of 7 different Y haplogroups mapping to GRCh38 for a 1000 Genomes Project variant for HG02982 (A0): Deletion nearby DDX3Y. Observed only in A0.

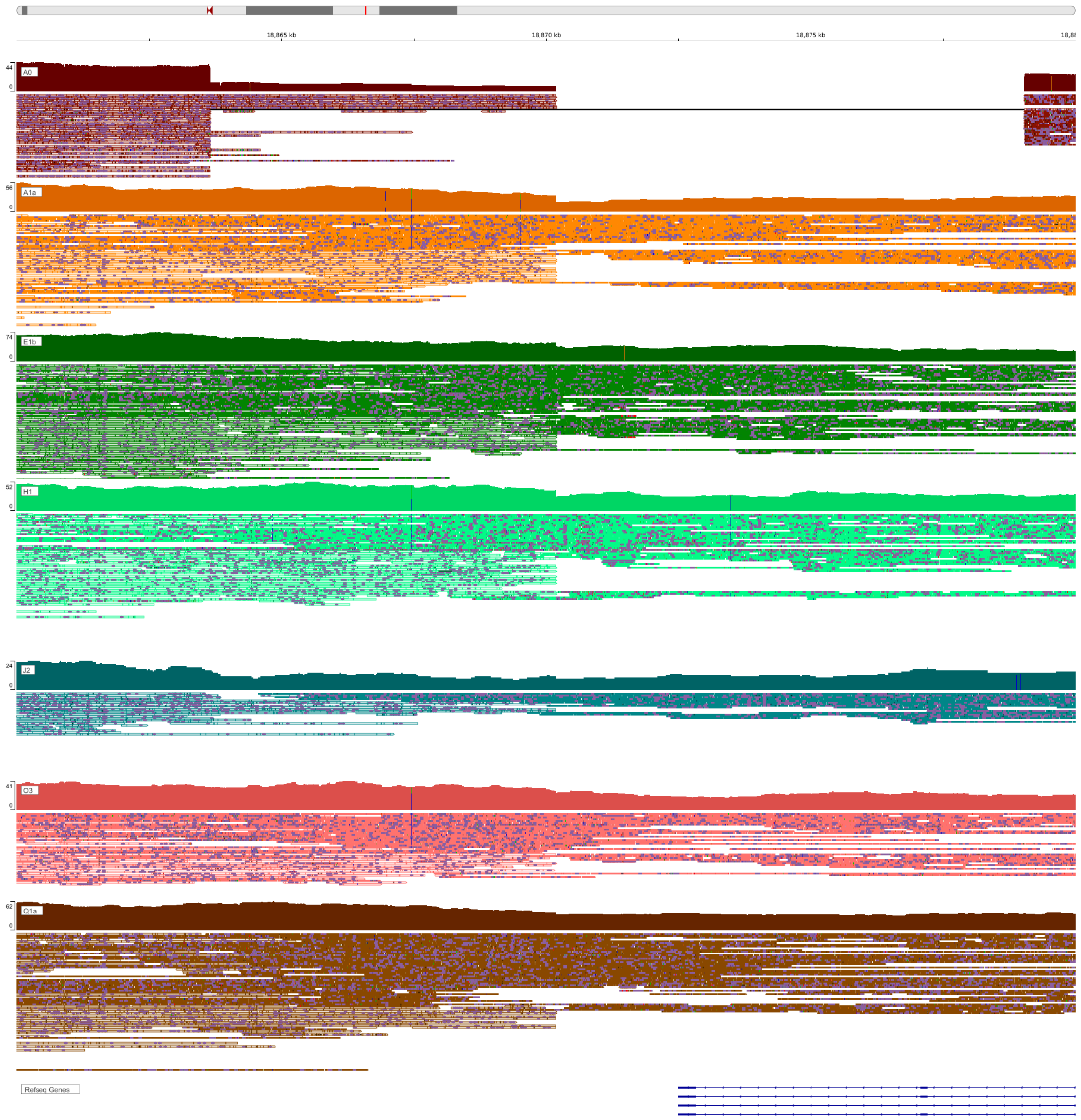


**Supplementary Figure 11.** IGV-screenshot showing ONT reads of 7 different Y haplogroups mapping to GRCh38 for a 1000 Genomes Project variant for HG02982 (A0): The deletion overlaps with the TTTY14 locus, nearby a segmental duplication.

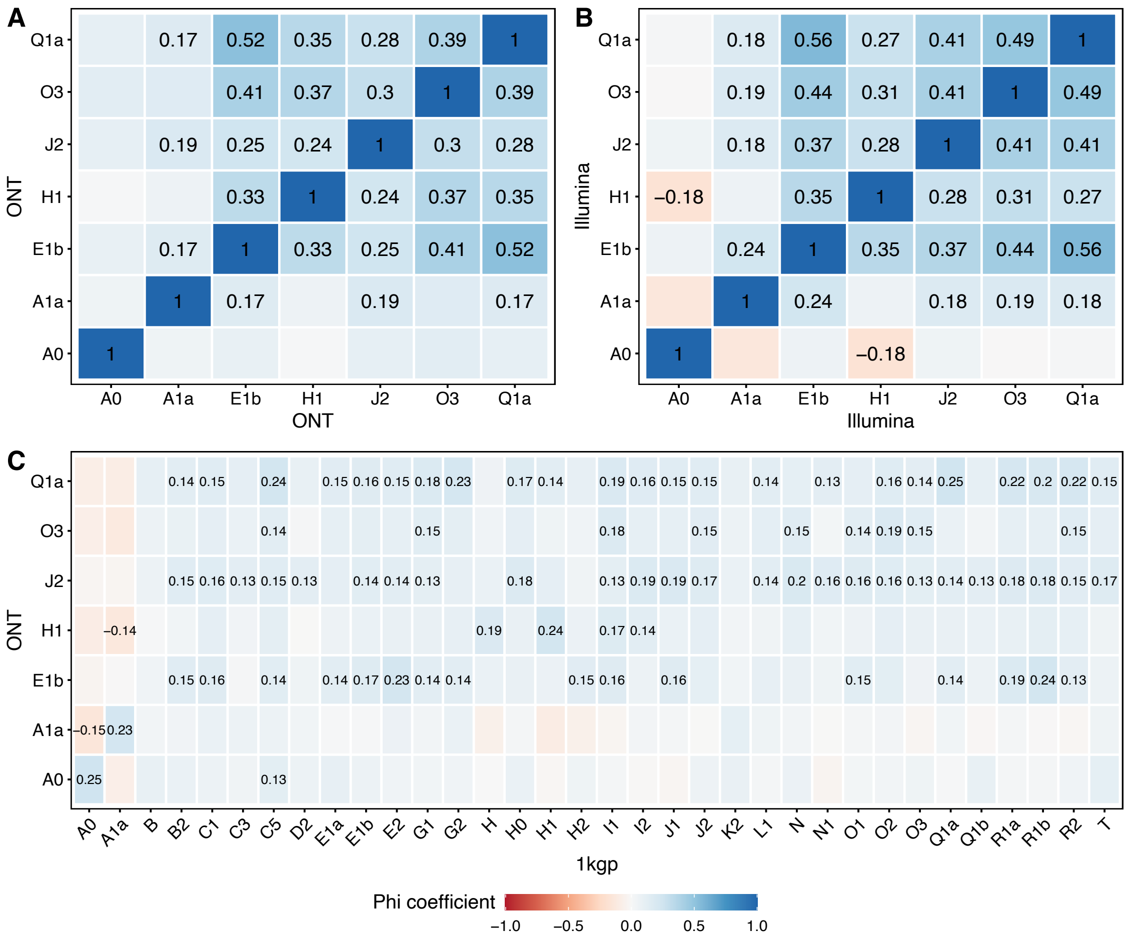


**Supplementary Figure 12.** Correlations of structural variant genotypes. (A) Correlations of genotypes obtained with *Sniffles v2* on ONT data. (B) Correlations of genotypes obtained using *graphtyper* on Illumina data. (C) Correlation of genotypes obtained using *graphtyper* on 1kgp short read data with the ONT data. Structural variants found in frequencies > 0.2 for the 1kgp data were considered as present and lower frequencies as absent. Only correlation values that are statistically significant after multiple testing correction are shown.

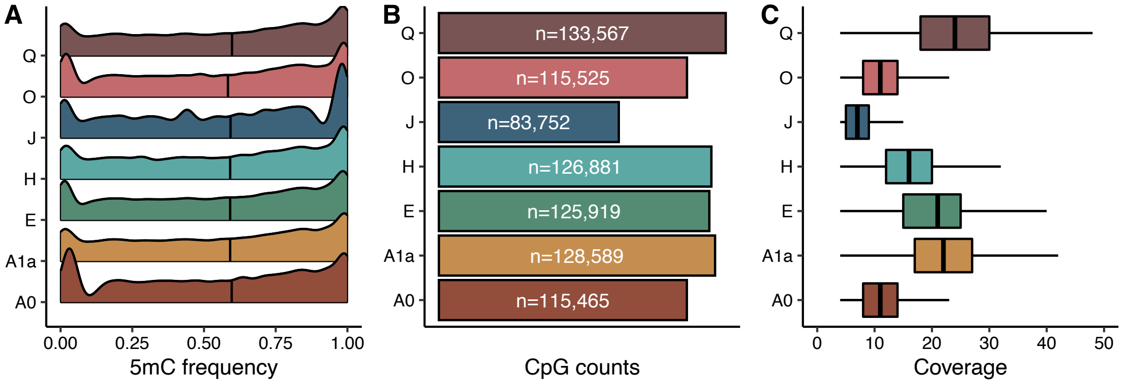


**Supplementary Figure 13.** (A) 5mC frequency levels, (B) Number of CpGs and (C) Coverage for each sample after quantile normalization (minimum coverage considered is 4x). Vertical black lines in (A) represent median 5mC frequency values.

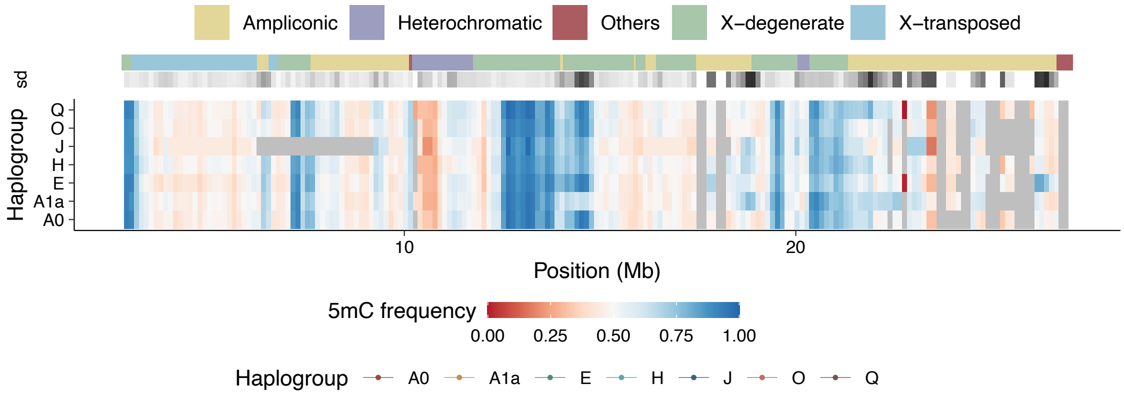


**Supplementary Figure 14.** Consistency in the frequency of 5mC in the seven cell lines along the resolved MSY of the hg38 is shown as tiles. The methylation levels are calculated as the median 5mC frequency value in 250kb sliding windows for each cell line. Each window requires a minimum of 10 CpG sites with methylation information. The sequence classes and the standard deviation of the methylation levels across cell lines are also shown. The standard deviation of the 5mC frequency is represented in a white-to-black scale, in which a darker color denotes a higher standard deviation value.

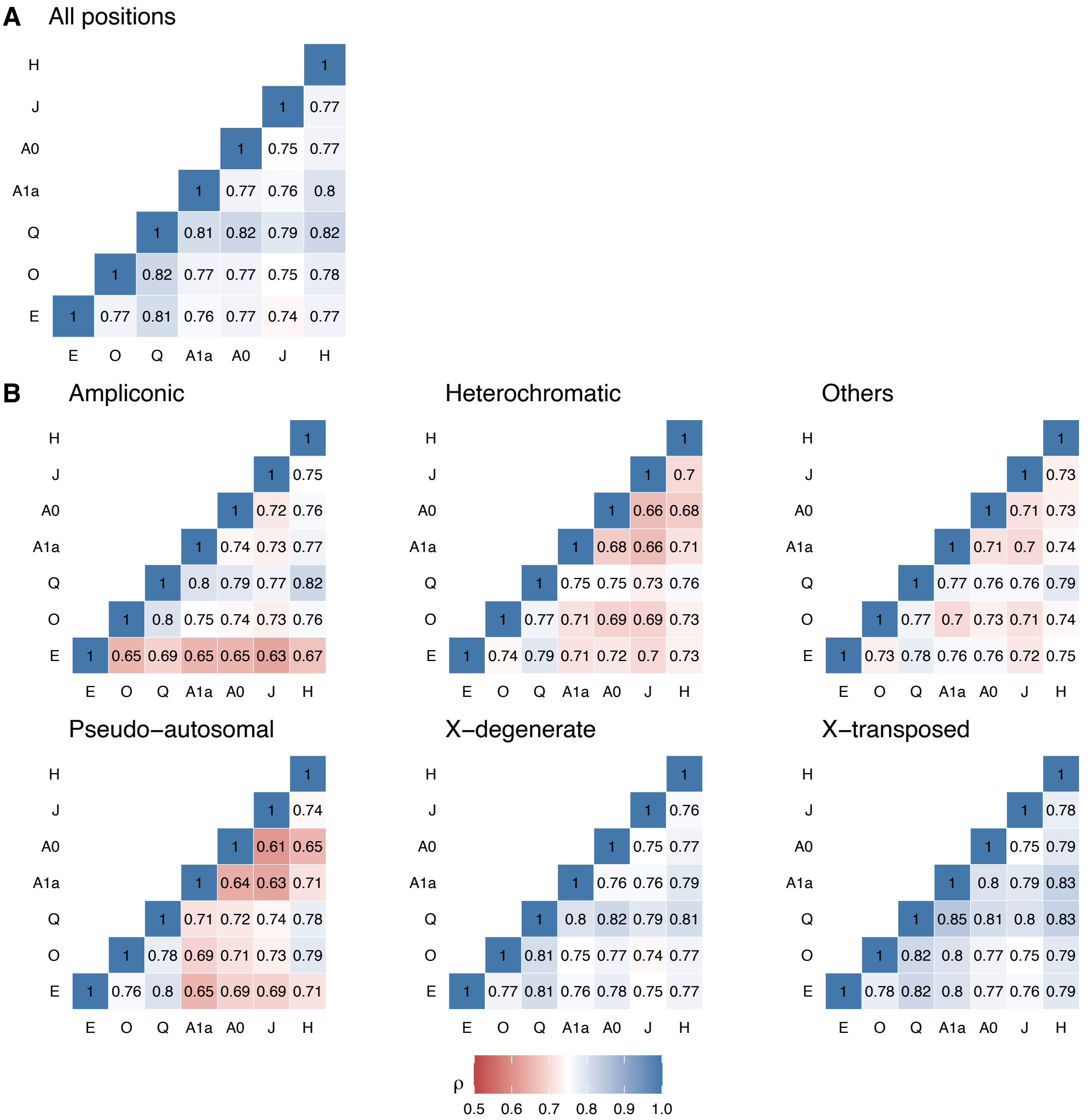


**Supplementary Figure 15.** High correlation of 5mC frequency in the seven cell lines calculated using the Spearman pair-wise correlation in (A) all CpG positions and in (B) sequence classes.

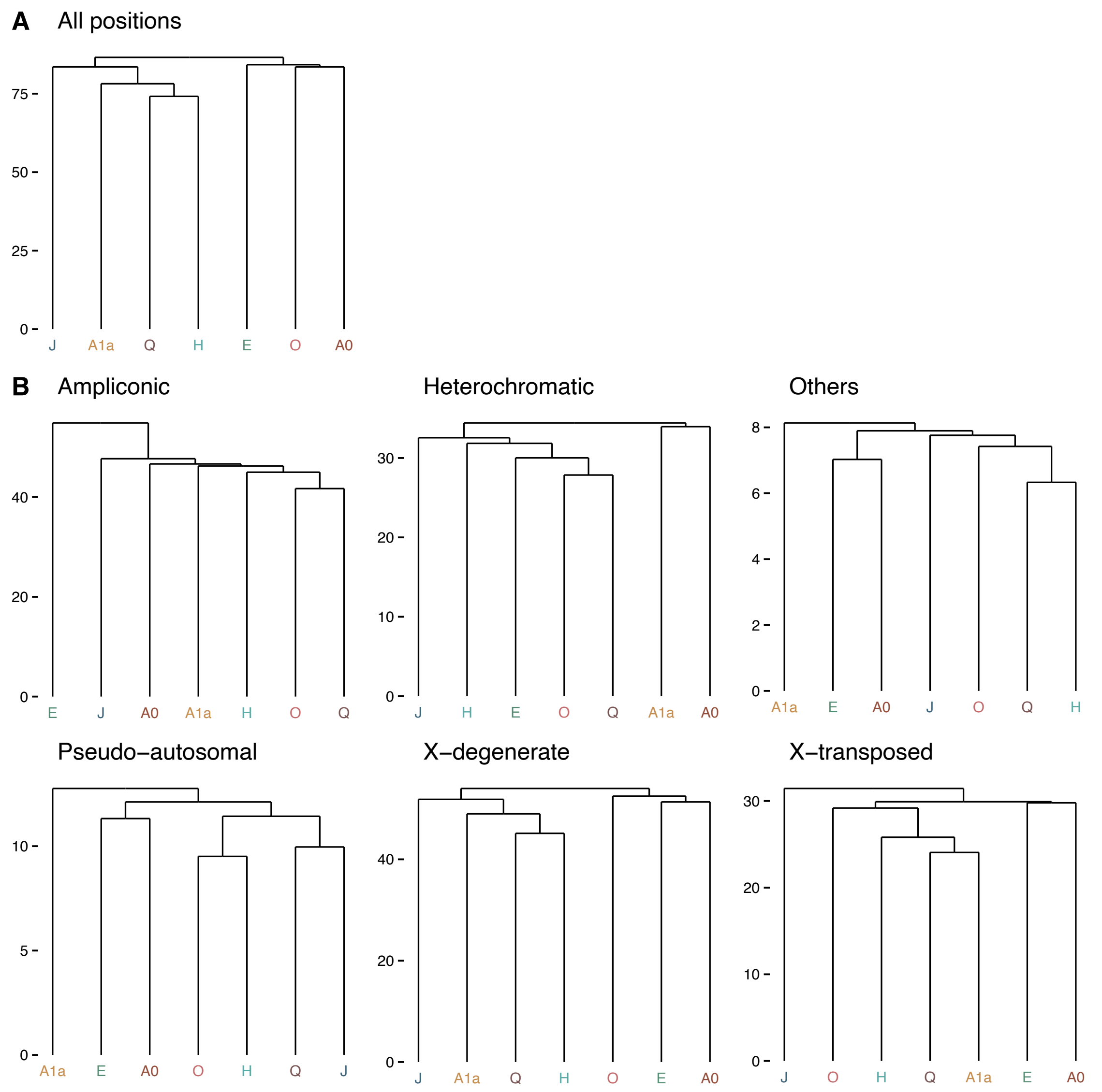


**Supplementary Figure 16.** Hierarchical cluster tree calculated using the Euclidean distance of 5mC frequency in the seven cell lines using (A) all CpG positions and (B) sequence classes.

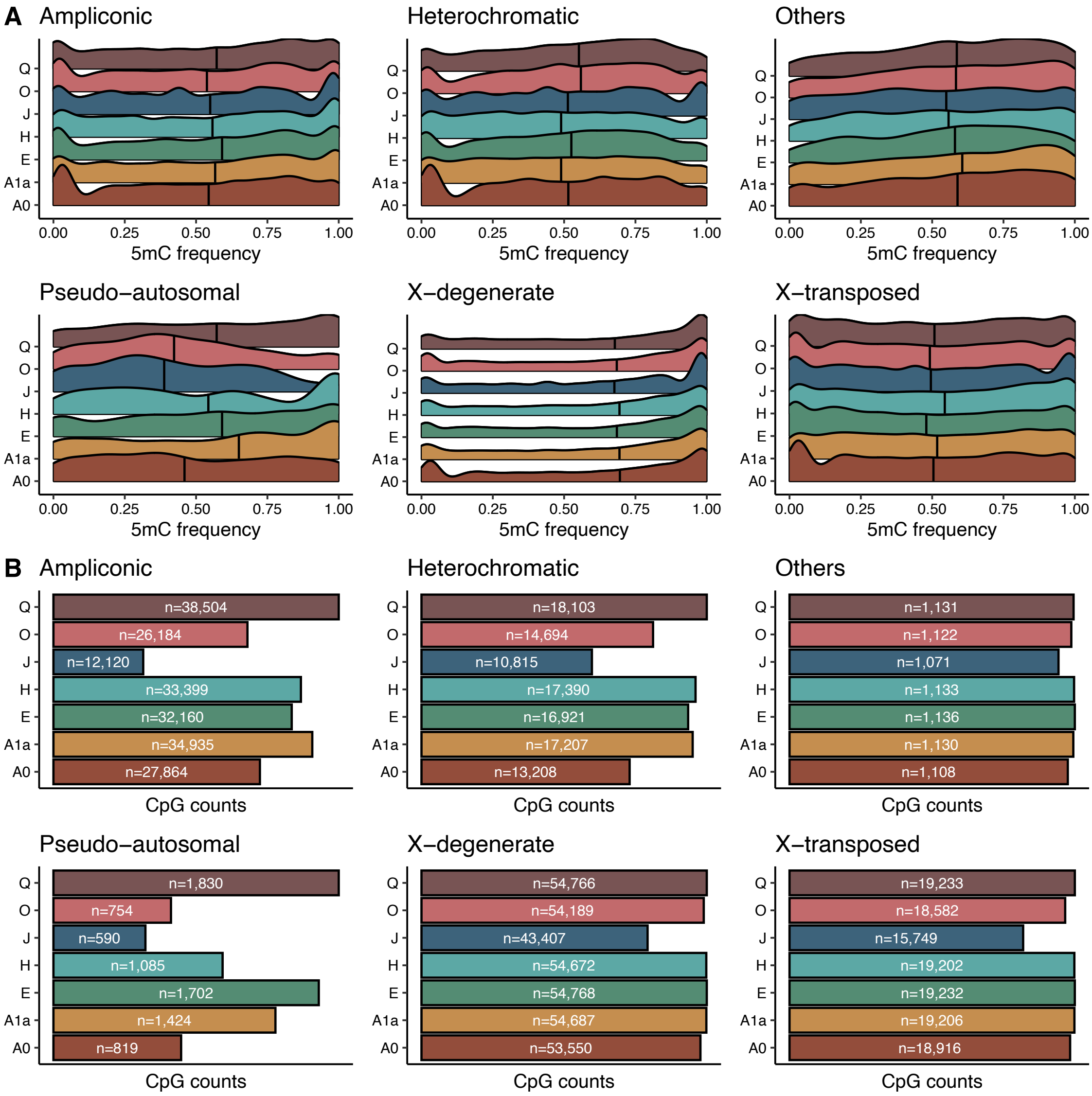


**Supplementary Figure 17.** Methylation profiles of each haplogroup per chromosome Y sequence classes. (A) 5mC frequency levels, (B) number of CpG in each sequence class used in (A). Vertical black lines in (A) represent 5mC frequency median values.

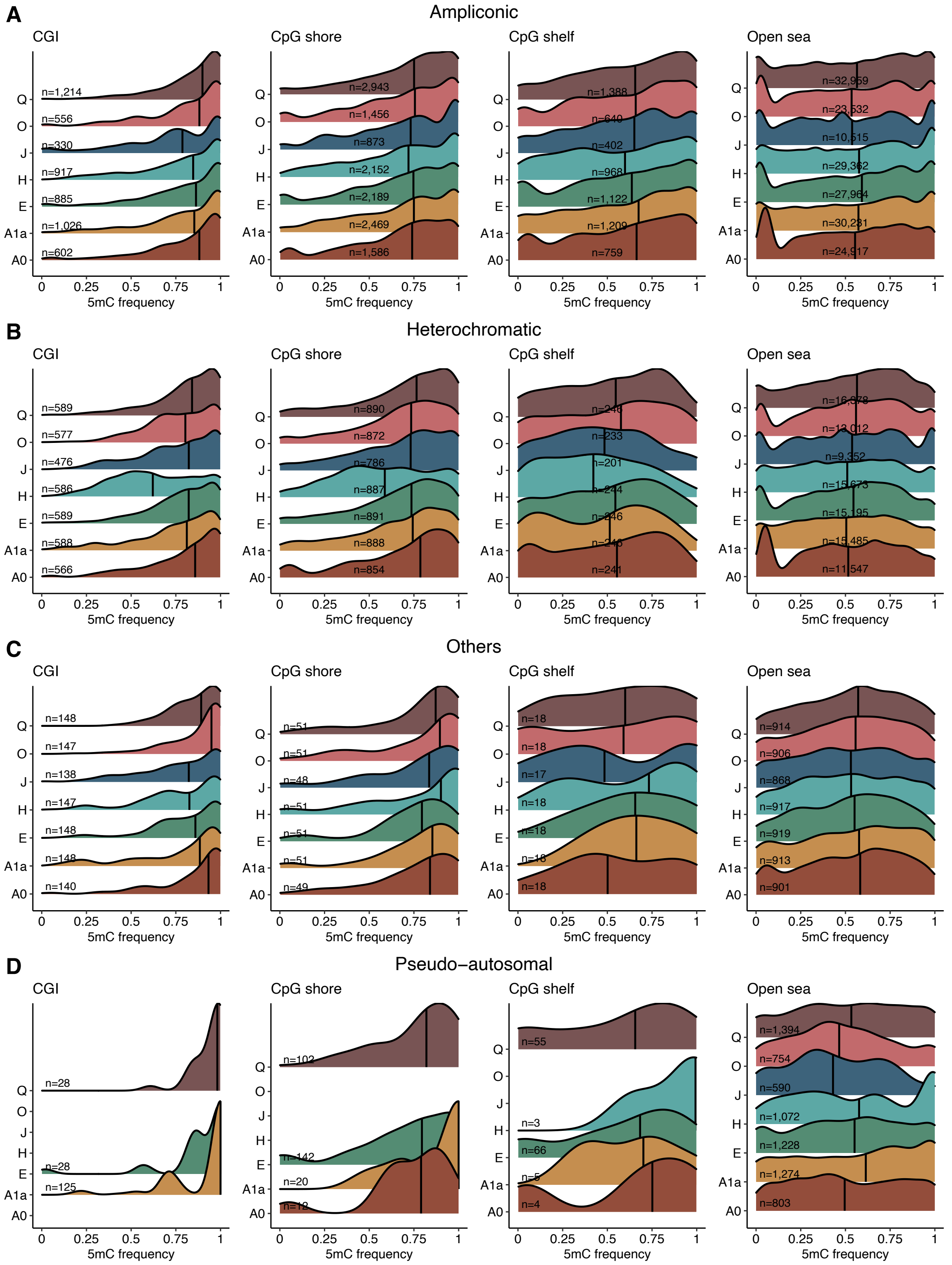


*(continues next page)*

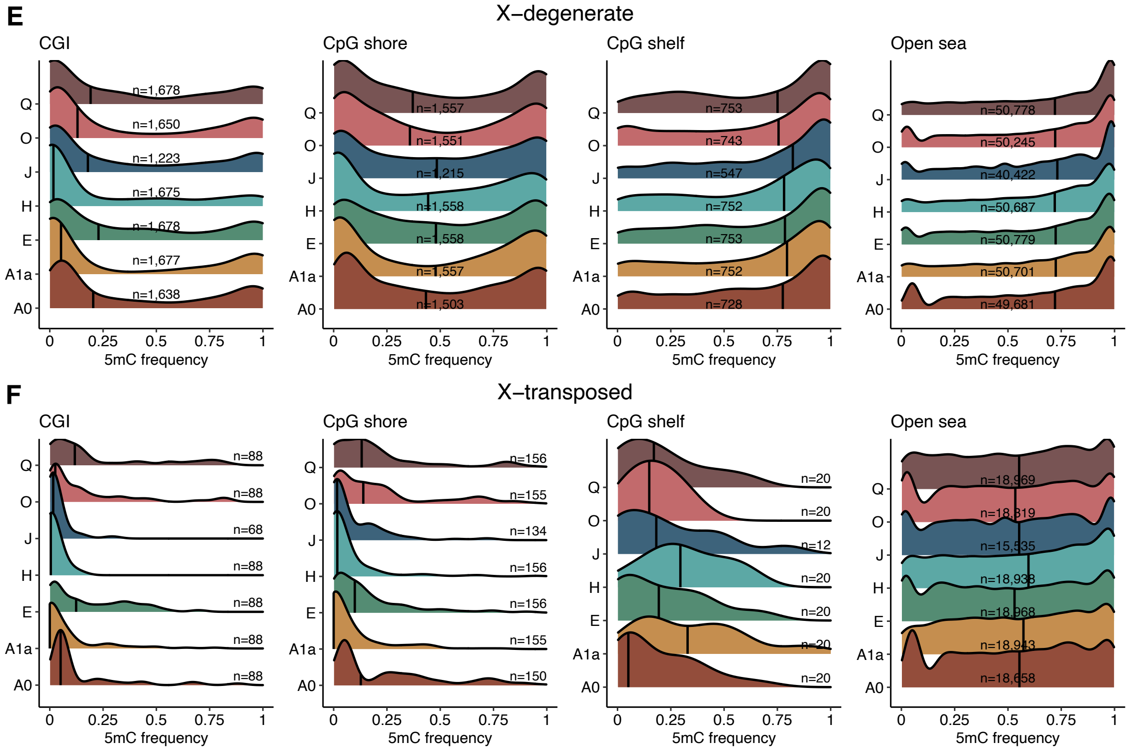


**Supplementary Figure 18.** Methylation profiles per chromosome Y sequence classes and CpG annotation: (A) ampliconic, (B) heterochromatin, (C) other regions, (D) PAR, (E) X-degenerate, and (F) X-transposed region. CpG annotations are mutually exclusive regions that comprise: CpG islands (CGI), CpG shores (up to 2kb away from the end of the CGI), CpG shelves (up to 2kb away from the end of the CpG shores), and inter-CGI or open sea regions (where all remaining CpG are allocated). The number of CpG in each annotation is shown within the plot. Vertical black lines represent median 5mC frequency values

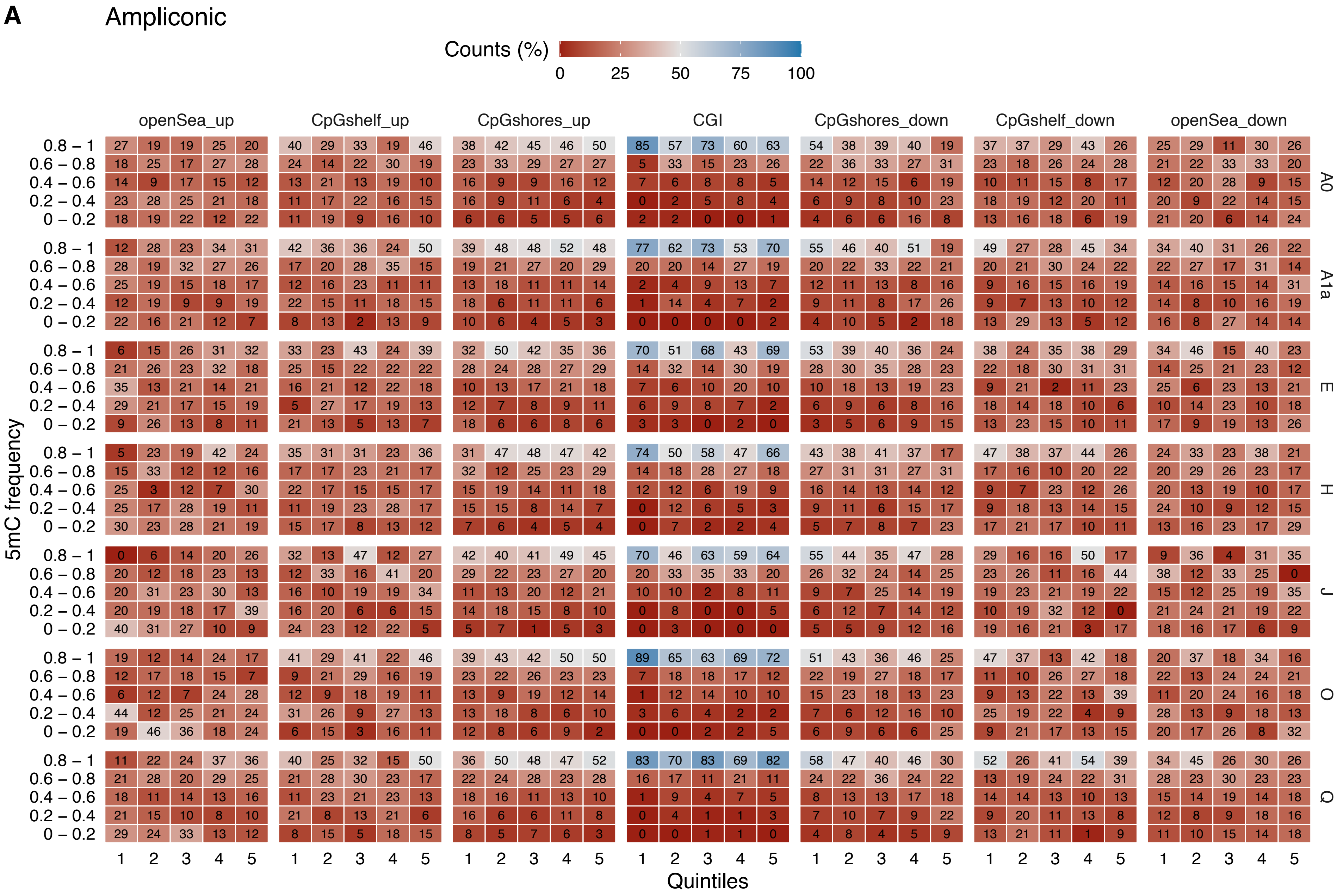
A
Ampliconic
Counts (%)
0
25
50
75
100
openSea_up
CpGshelf_up
CpGshores_up
CGI
CpGshores_down
CpGshelf_down
openSea_down
0.8 – 1
0.6 – 0.8
0.4 – 0.6
0.2 – 0.4
0 – 0.2
A0
A1a
E
H
J
O
Q
5mC frequency
1
2
3
4
5
Quintiles

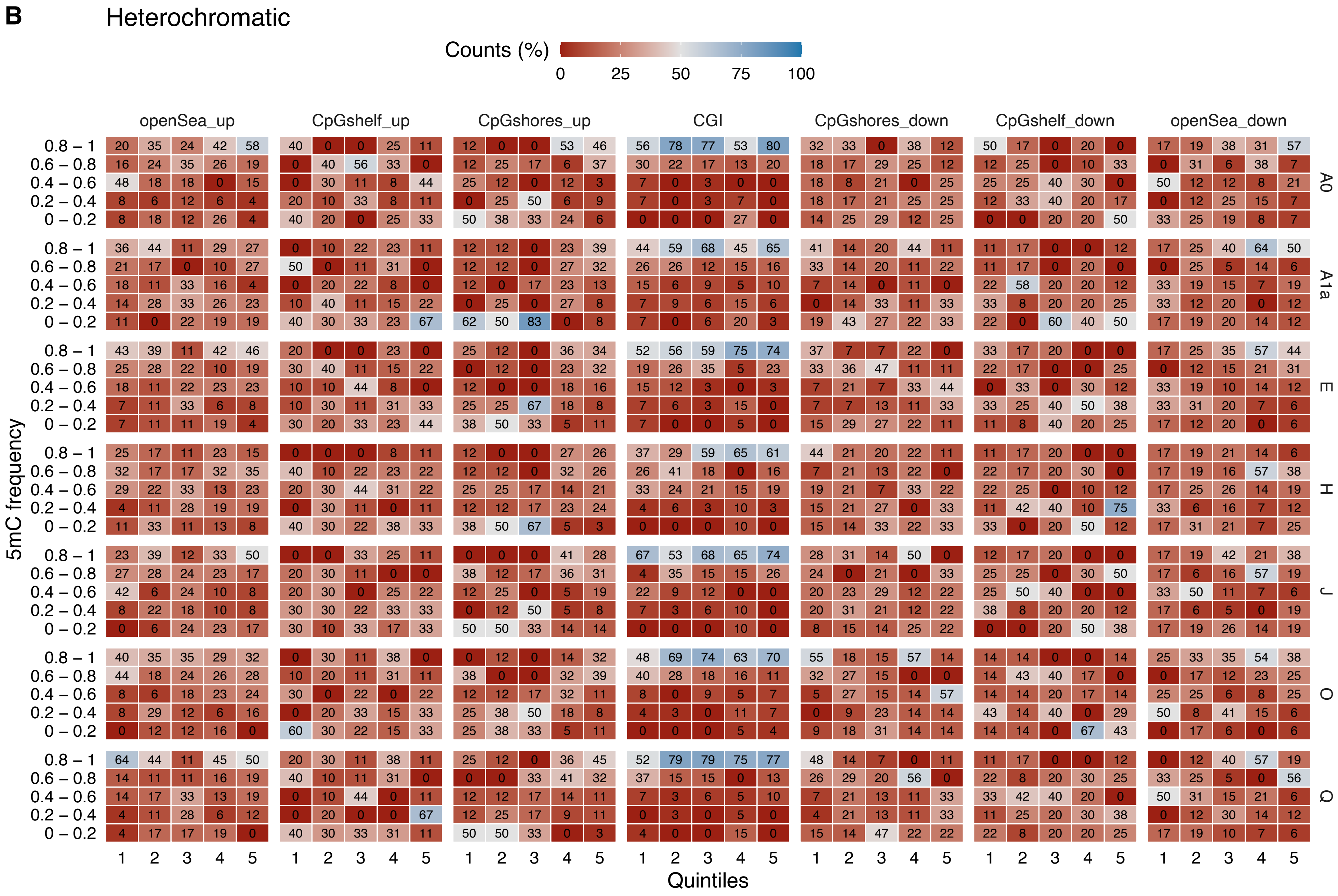
B
Heterochromatic
Counts (%)
0
25
50
75
100
openSea_up
CpGshelf_up
CpGshores_up
CGI
CpGshores_down
CpGshelf_down
openSea_down
0.8 – 1
0.6 – 0.8
0.4 – 0.6
0.2 – 0.4
0 – 0.2
5mC frequency
A0
A1a
E
H
J
O
Q
1 2 3 4 5
Quintiles

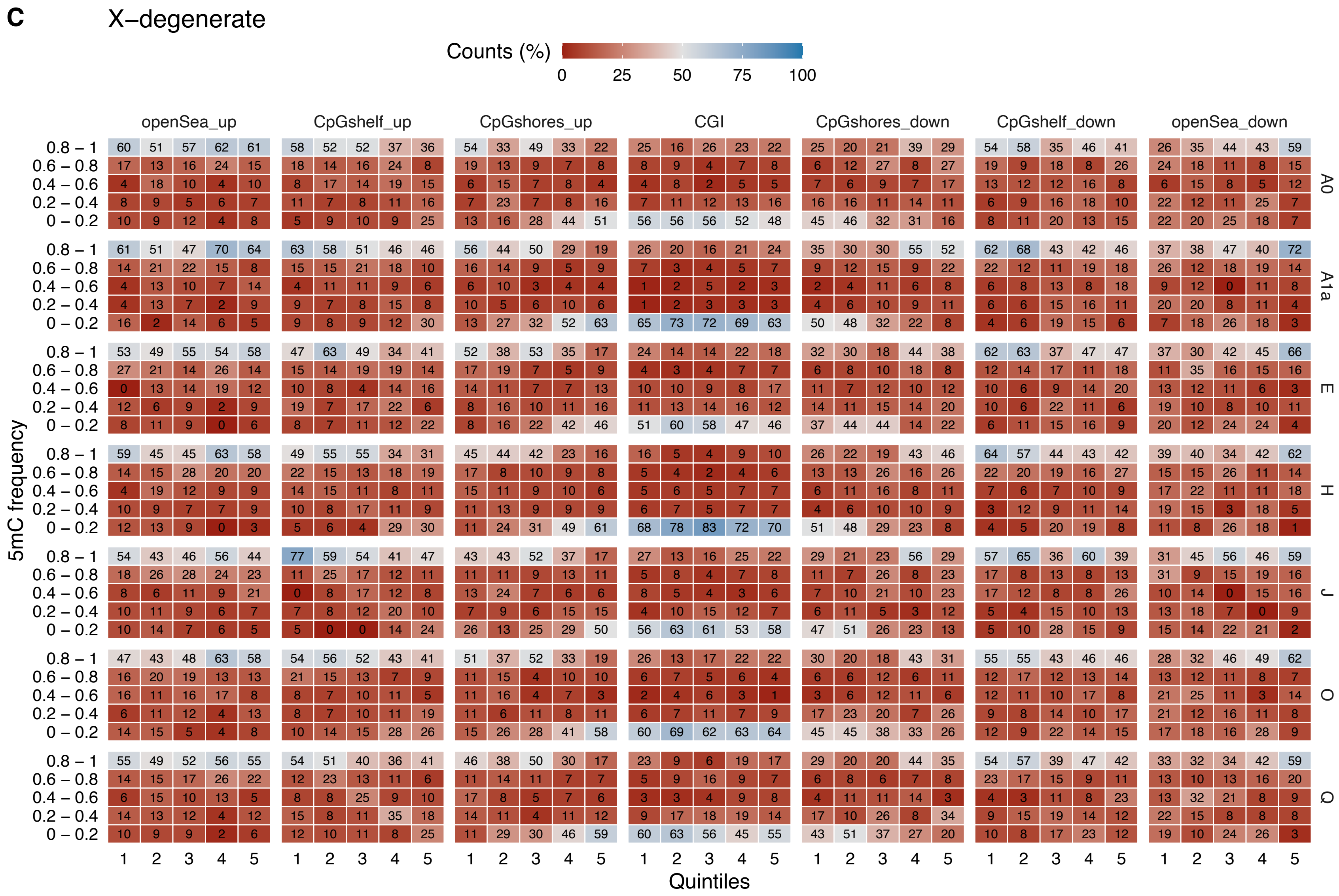


**Supplementary Figure 19.** Methylation frequency dispersion per CpG annotation in three sequence classes: (A) ampliconic, (B) heterochromatin and (C) X-degenerate region. For each CpG annotation, tiles show the proportion of CpG with a specific 5mC frequency. For that, CGI were only considered if they presented the full landscape (CpG shores, shelves and open sea) upstream and downstream of the CGI. Open sea was restricted to the 2kb regions closest to the CpG shelves. To account for the progression of methylation in each annotation, each CpG annotation was divided in 5 bins of equal length (quintiles). PAR, x-transposed and other regions were not represented as they did not have enough CpG sites.

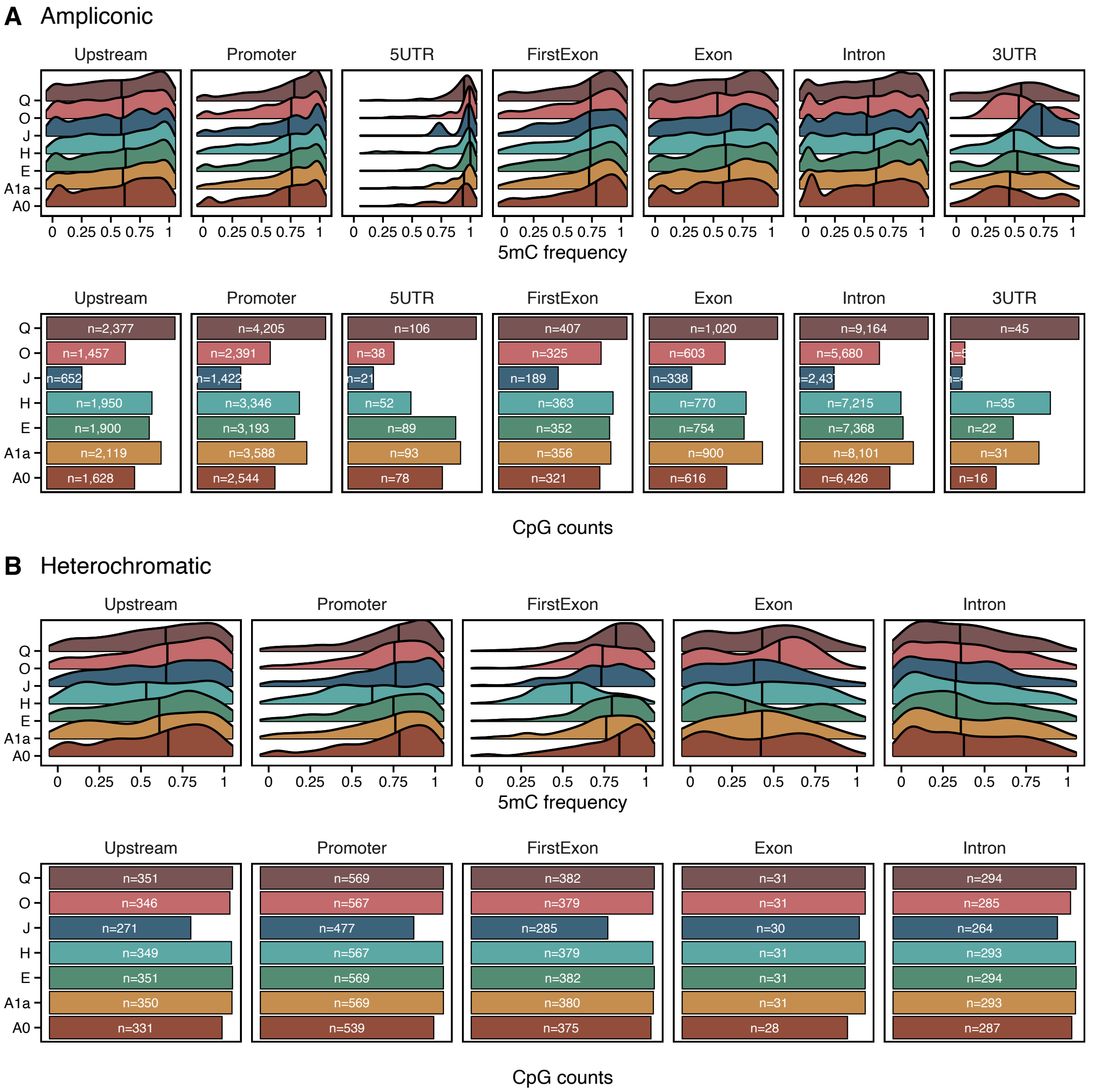


*(continues next page)*

**C** X–degenerate

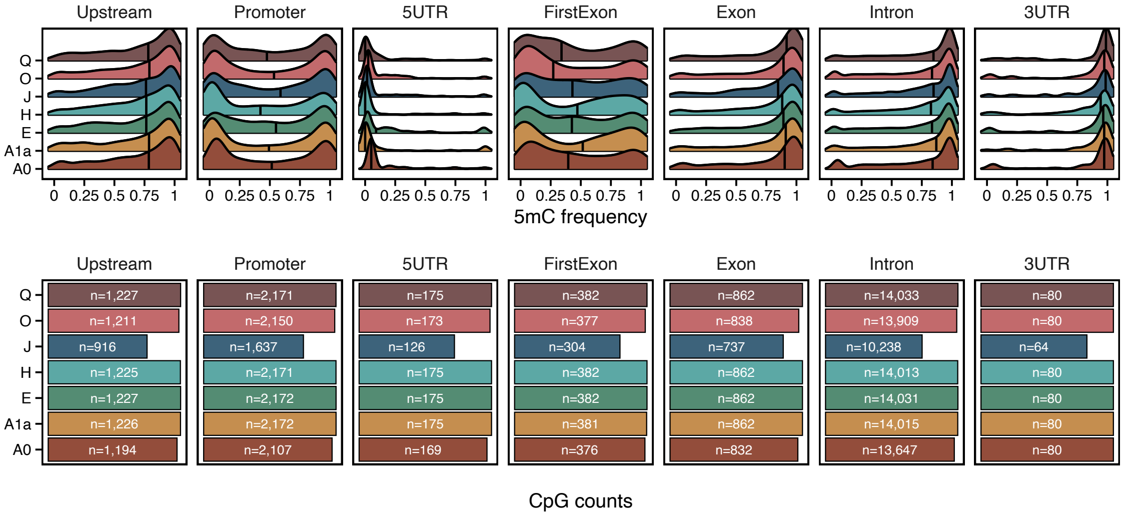


**D** X–transposed

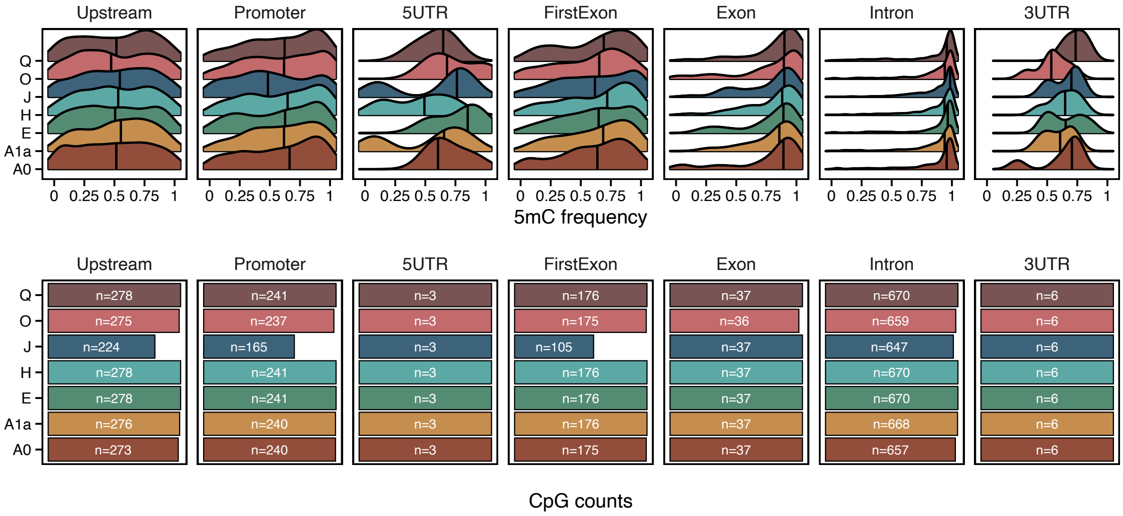


**Supplementary Figure 20.** Methylation profiles per chromosome Y sequence classes and CpG annotation: (A) ampliconic, (B) heterochromatin, (C) x-degenerate and (D) x-transposed region. Gene annotations are mutually exclusive regions that comprise the upstream region (up to 3kb from the end of the promoter), promoter (2kb upstream of TSS), 5'UTR, first exon, exons, introns, 3'UTR. For the overlapping regions, the priority set was the following: promoters, UTRs (5', 3'), first exon, other exons, all introns, and the upstream region. The number of CpG for each gene annotation is shown in the lower plot of each panel. Vertical black lines in each top plot represent median 5mC frequency values. Only chromosome Y sequence classes with most CpG sites were represented.

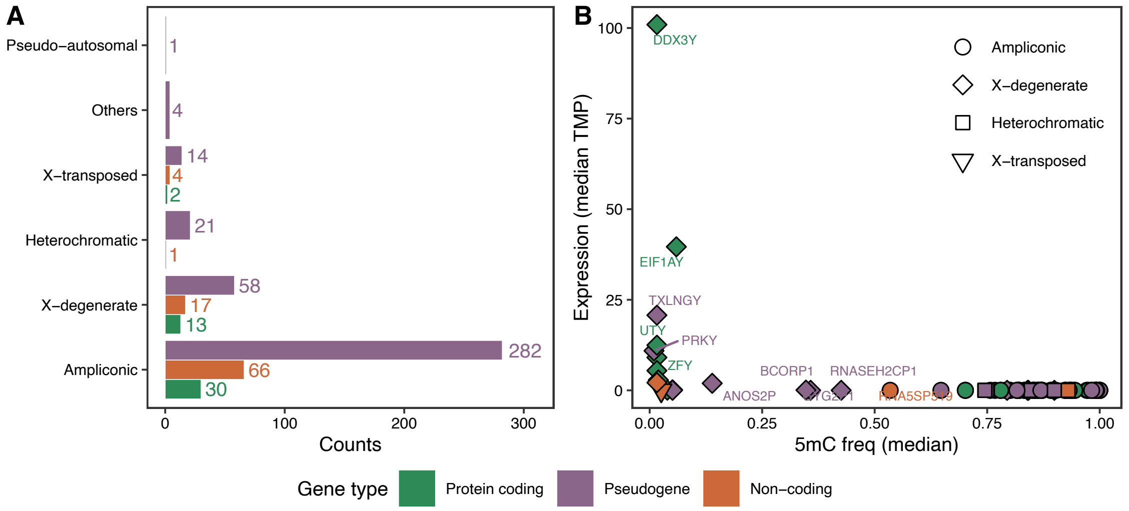


**Supplementary Figure 21.** (A) Number of genes per sequence class in the chromosome Y. The two most gene-dense regions are the ampliconic and X-degenerate regions. (B) Negative association between CGI methylation and gene expression. Each gene is associated with one upstream CGI and the median 5mC frequency is calculated for the 7 samples. Gene expression obtained is from GTEx data (v8) on LCLs derived from male samples and represented as median TMP.

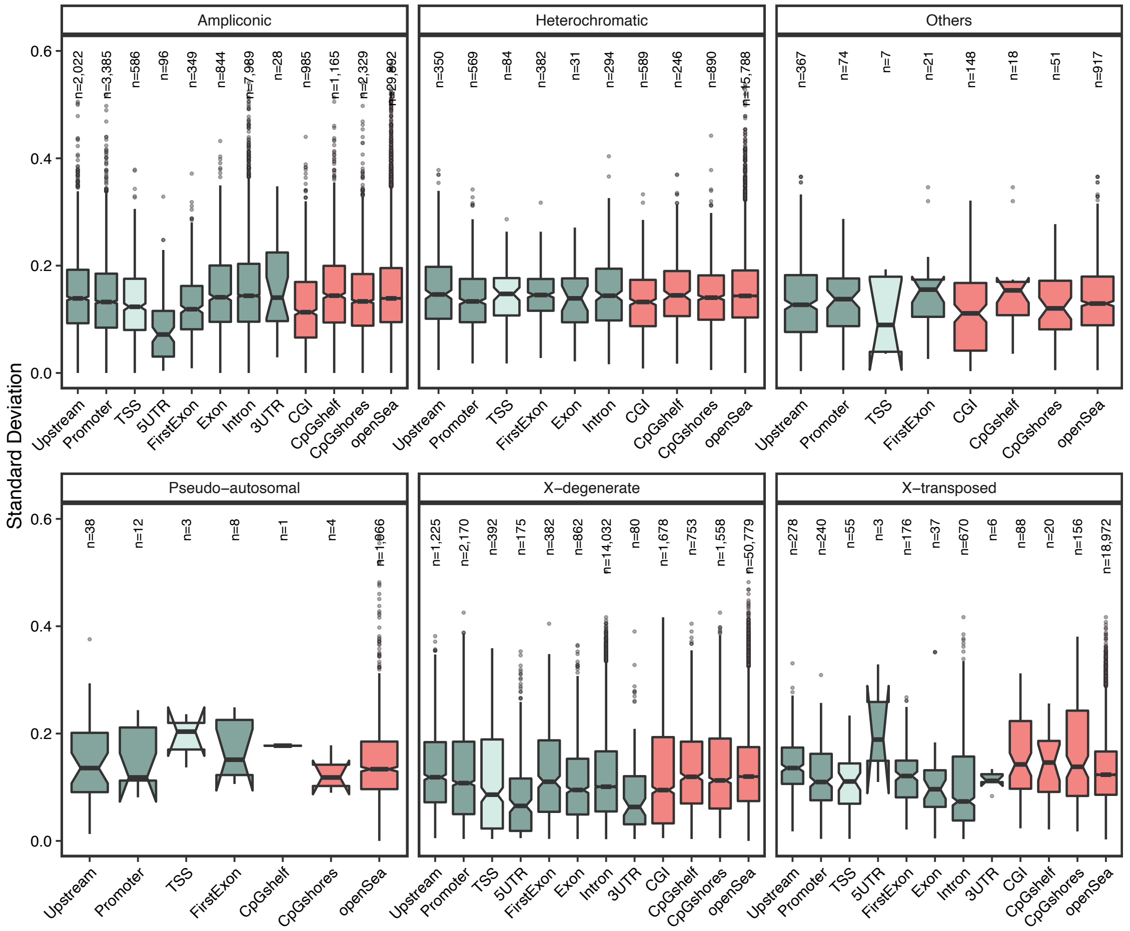


**Supplementary Figure 22.** CpG 5mC frequency dispersion between cell lines across sequence class and annotation. Dispersion is measured as the standard deviation of 5mC frequencies in CpG sites with methylation information for 3 or more cell lines. Gene annotation is coloured in blue while CpG annotation is coloured in salmon. Gene annotations are mutually exclusive regions as described in Supplementary Figure 6 in all instances but in TSS, which is a subset of the promoter region comprising the 200bp immediately upstream the TSS. The number of CpG sites used in each category is shown.

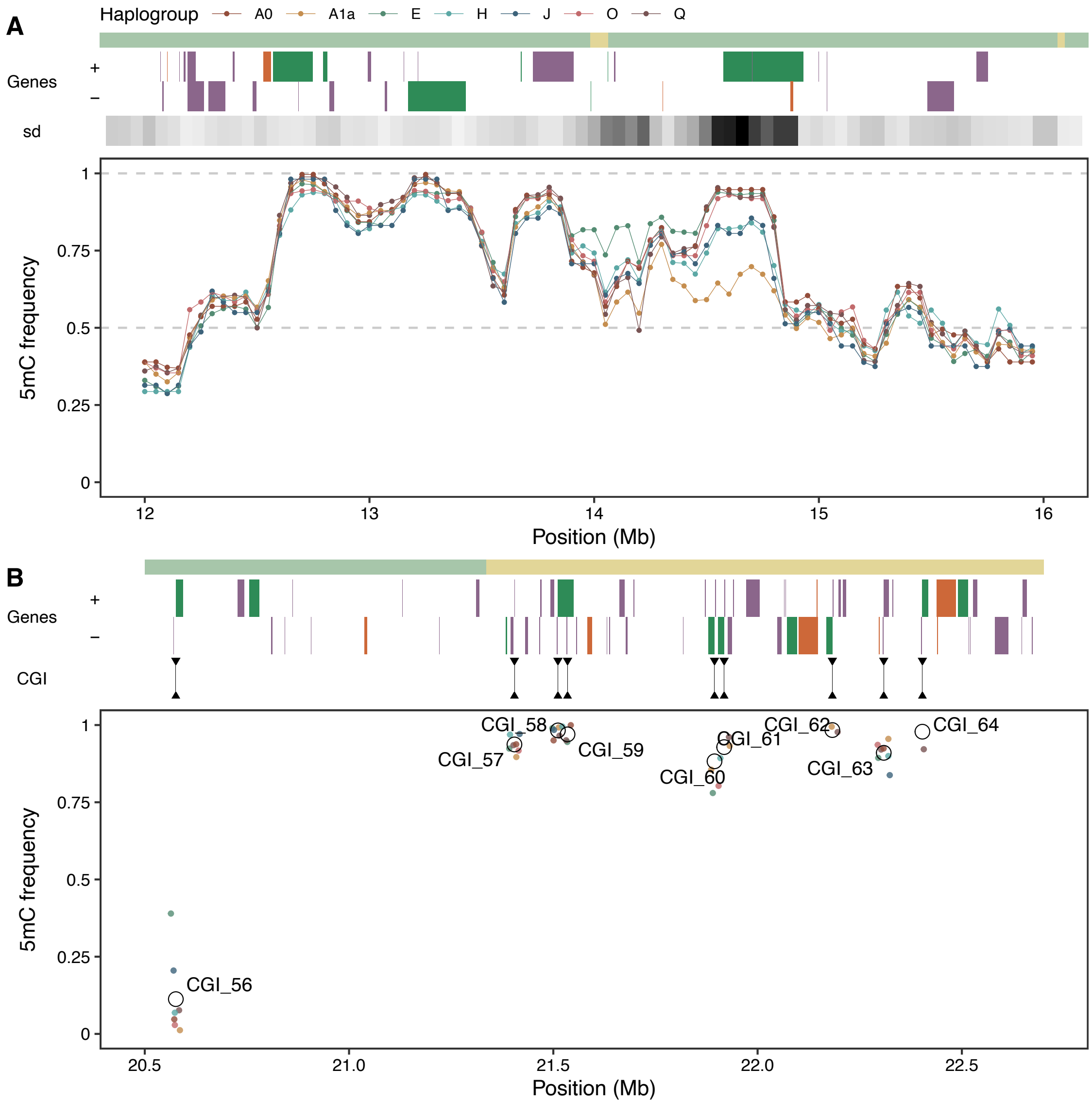


**Supplementary Figure 23.** (A) Frequency of 5mC in the seven cell lines along a specific region of the resolved MSY of the hg38: coordinates from 12 to 16 Mb. The methylation levels are calculated as the median 5mC frequency value in 100kb sliding windows for each cell line. The sequence classes, the genes annotated and the standard deviation of the methylation levels across cell lines are also shown. The standard deviation of the 5mC frequency is represented in a white-to-black scale, in which a darker color denotes a higher standard deviation value. The greatest dispersion region which has a higher standard deviation coincides with the protein-coding gene *NLGN4Y* (14.5 to 14.9 Mb). The cell line derived from haplogroup A1a shows lower 5mC values in this region. (B) Methylation frequencies in different CpG islands (CGI) in a specific region of the resolved MSY of the hg38: coordinates from 20.5 to 22.5 Mb. Empty circles denote the mean 5mC frequency per CGI whereas smaller colored points indicate the individual value in each cell line. CGI 56, which shows overall low 5mC frequency values is consistent with the regulation of a neighboring protein-coding gene in the positive strand, *EIF1AY*. On the other hand, CGI in the ampliconic region shows overall high 5mC values, also consistent with the silencing of most genes present in this region, of which the protein-coding genes belong to the *RBMY* gene family.
